# Arthropod predator nutrient content changes with crop sowing period with implications for biocontrol

**DOI:** 10.64898/2025.12.12.693942

**Authors:** Rosy Christopher, Rebecca Wright, Fredric M. Windsor, Jordan P. Cuff

## Abstract

Conservation biocontrol is crucial for crop pest management, but the impact of agricultural management practices on natural enemies of crop pests is poorly characterised. The timing of crop sowing could impact arthropod communities by altering basal resource availability and quality, manifesting in altered nutrient availability and cascading impacts to natural enemy fitness and function. This study aims to understand these potential modifiers of conservation biocontrol, by investigating how the timing of crop sowing impacts arthropod predator nutrition and whether associated differences relate to changes in prey availability. Arthropod predators and their prey were collected from adjacent plots of winter- and spring-sown wheat. The biochemical macronutrient (carbohydrate, lipid and protein) content of predators was determined by colorimetric assays and compared between winter-and spring-sown crops using multivariate models. Predator nutrient contents were compared to predator-prey interactions inferred using null network models based on both recent and current prey availability. The nutrient contents of arthropod predators differed significantly between winter- and spring-sown wheat, and the prey available to predators similarly differed, but neither recent nor current prey availability explained differences in predator nutrition. These findings may indicate that prey quality, rather than identity, changes with the developmental stages of crops sown at different times. This has important implications for the spillover of natural enemies of crop pests between crops sown at different times. This is especially true following harvest of winter-sown crops, which may disrupt biocontrol by altering the nutrients available to and sought by natural enemies of crop pests.

## Introduction

Conservation biocontrol, the promotion of natural enemies of crop pests through land management, can contribute substantially to pest suppression (Symondson et al., 2002; Tschumi et al., 2018). Whilst it effectively suppresses disease vectors and crop herbivores (Michalko et al., 2019; Moore et al., 2010), inefficiencies in conservation biocontrol arise through the predation of fellow natural enemies of crop pests and other beneficial invertebrates, such as pollinators (Hodge, 1999; Rosenheim et al., 1995; Tschumi et al., 2018). To optimise the biocontrol activity of natural enemies of crop pests, we must understand how agricultural management decisions affect natural enemy populations and their interactions with pests, including the myriad changes that arable land progresses through across the annual cycle (Cuff, Gajski, et al., 2024). From harvest practices (Cuff et al., 2021; Opatovsky & Lubin, 2012), through tillage (Heimbach & Garbe, 1996; Jowett et al., 2021), to the timing of sowing (Douglas et al., 2010), management decisions can have profound implications for conservation biocontrol (Cuff, Gajski, et al., 2024).

The season within which crops are sown can greatly impact their management and productivity; for example, spring-sown crops can see improved disease delay relative to winter-sown crops (Landa et al., 2004). As a result, land managers have developed a diverse range of crop management strategies and cycles. Such variations in crop management will also manifest in differences in invertebrate community structure and function due to cascading impacts arising from differences in crop developmental stages (Douglas et al., 2010; Hsu et al., 2021). Furthermore, it has the potential to impact the interactions of pests and their predators; for example, when crops sown at different times co-occur in an agricultural landscape, spillover of invertebrates and their interactions from one crop to another presents a significant opportunity for the spread of both pests and predators, especially when earlier-sown crops are harvested (Cuff, Gajski, et al., 2024; Jaworski et al., 2023). By modifying invertebrate community structure (Douglas et al., 2010), agricultural management decisions like crop sowing timing can also indirectly impact the nutrients available to and consumed by natural enemies of crop pests by altering crop development and senescence phenology. As crops senesce in their later developmental stages, nutrients are remobilised away from leaf tissues (Gregersen et al., 2008), causing foliar herbivores and their predators to migrate or die (Watt, 1979).

Nutrients are the fundamental currency of trophic interactions, influencing population dynamics, physiology and species distributions, and driving biocontrol and other ecosystem services (Cuff, Evans, et al., 2024; Wilder et al., 2025). Disruption to nutrient acquisition can cascade through consumers (Hedlund et al., 2004), reducing energy transfer between trophic groups (Rosenblatt, 2018), ultimately disrupting ecosystem service provision. Generalist predators redress nutritional deficits that arise through such perturbations by engaging in nutrient-specific foraging, the selection of prey based on nutrient content, to optimise their nutrient intake, and therefore fitness (Jensen et al., 2012; Mayntz et al., 2005). Nutritional differences between prey are therefore important for the choices predators make, the trophic interactions they engage in and the ecosystem services they provide, and are likely to be sensitive to our land management practices. By understanding how nutritional dynamics drive ecological processes and are driven by our actions, we can optimise our land management to promote the provision of ecosystem services, including conservation biocontrol (Cuff, Tercel, Vaughan, et al., 2024).

Here, we investigated how crop sowing season affects the nutrient content of arthropod predators and whether this is driven by alterations to the prey community available to them. We collected arthropod predators and their prey from winter- and spring-sown wheat, and determined predator biochemical macronutrient (carbohydrate, lipid and protein) content. Using multivariate modelling and null network inference, we tested the following hypotheses: (i) invertebrate prey availability will differ between winter- and spring-sown crops; (ii) arthropod predator nutrient content will differ between winter- and spring-sown crops; (iii) differences in predator nutrient content will relate to recent, but not current, prey availability. By elucidating the nutritional consequences of management decisions such as sowing time for arthropod predators, we can begin optimising management to safeguard the fitness of natural enemies of crop pests to sustain their suppression of crop pests.

## Methods

### Fieldwork and identification

Arthropods were collected from Cockle Park (Morpeth, UK; 55°13’00.8” N, 1°41’28.7” W) in July 2024. Transects were established in a field containing adjacent plots of spring-sown and winter-sown wheat. Six pairs of belt transects were established 20 m apart within each crop with 4 m between adjacent transects in the spring and winter sown crops. One pitfall trap for each pair of transects (i.e., three per sowing period) was filled with soapy water and deployed for 72 h. Pitfall trapping was conducted once nine days prior to individual arthropod collection (to represent past prey availability) and once in parallel with individual arthropod collection (to represent present prey availability). Invertebrates within these traps were filtered in-field using an Aeropress^Ⓡ^ and stored in 80 % ethanol for subsequent morphological identification. Within each transect, hand-searching was conducted for approximately 10 minutes and individual arthropods collected using a pooter.

Following collection, the arthropods were frozen onsite at -20 °C to humanely kill them and preserve their macronutrient contents for subsequent analysis. The samples were transported to the Molecular Diagnostics Facility at Newcastle University for morphological identification and nutritional analysis. All invertebrates were identified using a stereomicroscope and morphological keys (Barber, 2008; Dallimore & Shaw, 2013; Luff, 2007; Roberts, 2016; Tilling, 2014). Most invertebrates were identified to at least family level or finer resolution (e.g., genus and species), except where access to appropriate identification resources or damage to specimens limited identification. Individuals from each taxa were then enumerated to produce species by site abundance data.

### Macronutrient determination

Macronutrient analysis followed the Macronutrient Extraction and Determination from Invertebrates (MEDI) method (Cuff et al., 2021; Cuff and Wilder, 2021) which streamlines sulfo-phospho-vanillin, anthrone and Lowry colorimetric assays for lipid, carbohydrate and protein estimation, respectively. To extract macronutrients from whole arthropods, individually collected samples were placed in 2 mL tubes of a 96-well rack and heated until dry (∼ 1 h) in an oven at 60 °C. To each sample, 500 μl of 1:12 chloroform:methanol solution was added and left at room temperature for 24 h. After 24 h, 400 μl of the chloroform:methanol solution was removed and retained separately for lipid analysis. A further 500 μl of chloroform:methanol solution was added to the sample and incubated at room temperature for 24 h to remove any residual lipids. All chloroform:methanol solution was removed by pipetting, and any residue evaporated in an oven at 60 °C for ∼ 15 min.

To each dried sample, 500 μl 0.1 M NaOH was added, after which the samples were incubated at 80 °C for 30 min and at room temperature overnight (∼ 16 h). The samples were briefly centrifuged (∼ 2 min, 2,000 x g) to avoid taking solid material through to the subsequent assays. From each sample, 400 μl of the supernatant was transferred into a separate tube for protein and carbohydrate determination. All assays were carried out in 384-well plate format. For each assay, a stock standard dilution series of a known concentration was made with lard oil, glucose and bovine serum albumin (BSA) for lipid, carbohydrate and protein quantification, respectively. Stock solutions of 2 mg ml^−1^ glucose and BSA were made using distilled water and subsequently diluted in 0.1 M NaOH, whereas 2 mg ml^−1^ lard oil was made and diluted with 1:12 chloroform:methanol. Dilution series comprised 0-1 mg ml^−1^ in nine increments (0, 12.5, 62.5, 125, 250, 375, 500, 750, 1000 μg ml^−1^). A quarter of each 384-well assay plate (96 wells) was reserved for repeats of these standards.

### Lipid quantification

The sulfo-phospho-vanillin method was used to determine lipid content. The vanillin reagent was prepared by mixing 0.108 mg of vanillin with 18 μl of hot water and 72 μl 85 % phosphoric acid for every 100 μl reagent. From each sample, three repeats of 20 μl were added to a 384-well plate alongside standards and heated to 100 °C for 5 min to evaporate the chloroform:methanol carrier. To each well, 10 μl of concentrated sulfuric acid was added, mixed briefly on a plate shaker and incubated at room temperature for 15 min. Following this, 75 μl of vanillin reagent was added to each well, mixed briefly on a plate shaker and incubated at room temperature for 10 min. Absorbance at 490 nm for each was determined in a spectrophotometer.

### Carbohydrate quantification

The anthrone method was used to determine carbohydrate content. The anthrone reagent was prepared by mixing 100 μg of anthrone with 100 μl of concentrated sulfuric acid for every 100 μl of reagent. From each sample, three repeats of 20 μl were added to a 384-well plate alongside standards, mixed with 80 μl of the anthrone reagent and incubated at 92 °C for 10 min. The plate was incubated at room temperature for 5 min and absorbance at 620 nm for each well was determined in a spectrophotometer.

### Protein quantification

The Lowry method was used to determine protein content. From each sample, three repeats of 20 μl were added to a 384-well plate alongside standards, and 60 μl of modified Lowry reagent added to each well and mixed briefly on a plate shaker. The plate was incubated at room temperature for 10 min, and 6 μl of 1X Folin-Ciocalteu reagent added and mixed briefly on a plate shaker. The plate was incubated at room temperature for 30 min and absorbance at 750 nm for each well was determined in a spectrophotometer.

### Statistical analysis

All analysis was conducted in R version 4.5.2 (R Core Team, 2025) and all data processing used the ‘tidyverse’ package for reproducibility (Wickham et al., 2019). Data visualisations were generated using ggplot2 (Wickham, 2016).

The pitfall trapped invertebrate communities (herein considered representative of prey available to the hand-collected arthropod predators) were compared between winter and spring sown crops, pitfall trapping rounds and an interaction between the two using multivariate generalized linear models (MGLMs) with a Poisson error family and Monte Carlo resampling in the ‘mvabund’ package via the ‘manyglm’ function (Wang et al., 2012). These differences were visualised using non-metric multidimensional scaling with a Bray-Curtis dissimilarity matrix in the ‘vegan’ package (Oksanen et al., 2022). Macronutrient contents of arthropods were compared between taxa, winter- and spring-sown crops and the interaction between the two using multivariate linear models in the ‘mvabund’ package via the ‘manylm’ function (Wang et al., 2012).

Null network models were run to infer density-dependent interactions for each of the individually-collected arthropod predators based on prey availability and determine if prey availability explains differences in predator nutrient content. Using pitfall trapping prey availability data, separately for the past and present prey availability based on the two rounds of pitfall trapping, null networks were generated for each of the individually-collected predators included in the nutritional analyses above. These null network models assume that the trophic interactions of the predators were density-dependent (i.e., they interacted with the prey most available to them), providing a semi-realistic estimate of how prey availability would translate to trophic interactions.

To construct null networks, null diets were simulated using the ‘generate_null_net’ function in the ‘econullnetr’ package (Vaughan et al., 2018) using the prey availability data from the pitfall traps with random placeholder interaction data. To generate null networks, we required dummy interaction data. We simulated these dummy interactions for each individual arthropod predator by randomly allocating interactions with three of the available prey taxa (approximately reflecting the interaction richness of arable arthropod predators from previous work; Cuff et al., 2022). These dummy data were only required to run null network simulations and did not otherwise feature in any downstream analyses, with the exception of the fact that the node degree is invariable (i.e., a mean of three interactions per predator across all simulations). Using the ‘generate_null_net_indiv’ function (Cuff et al., 2023), just the simulation data were extracted irrespective of the identities of the random interaction data input. Null networks were visually constructed using ‘igraph’ (Csardi & Nepusz, 2006) and plotted using ‘ggnetwork’ (Briatte, 2021). Null network-inferred predator-prey interactions based on both past and present prey availability were compared against predator nutrient content using redundancy analysis (RDA) and model outputs analysed using analysis of variance (ANOVA) of the overall model, terms within the model and model axes.

## Results

From the six transects, nutritional data were generated for 98 hand-collected predatory arthropods, 52 from the winter-sown wheat, and 46 from the spring-sown wheat. These belonged to five families: Carabidae (n = 86), Coccinellidae (2), Henicopidae (3), Linyphiidae (3) and Staphylinidae (4). The pitfall traps collected 668 individual invertebrates across 28 taxa resolved to at least family level, considered representative of prey available to the above predatory arthropods. Available prey communities differed significantly between crops sown in winter and spring (MGLM: Dev = 213.55, df = 10, p = 0.001; Figure 2), between samples taken in parallel to hand-collected predators and nine days prior (MGLM: Dev = 133.33, df = 9, p = 0.001) and the interaction between sowing period and sampling time (MGLM: Dev = 42.98, df = 8, p = 0.001). Specifically, the abundances of brachyceran flies (MGLM: Dev = 20.794, df = 10, p = 0.001), carabid beetles (MGLM: Dev = 28.456, df = 10, p = 0.001), ichneumonid wasps (MGLM: Dev = 8.662, df = 10, p = 0.046), linyphiid spiders (MGLM: Dev = 23.599, df = 10, p = 0.001), platygastrid wasps (MGLM: Dev = 83.939, df = 10, p = 0.001) and stratiomyid flies (MGLM: Dev = 9.704, df = 10, p = 0.034) differed between crops sown in winter and spring. Similarly, the abundances of brachyceran flies (MGLM: Dev = 20.794, df = 9, p = 0.001), carabid beetles (MGLM: Dev = 9.272, df = 9, p = 0.044), linyphiid spiders (MGLM: Dev = 31.662, df = 9, p = 0.001) and stratiomyid flies (MGLM: Dev = 9.704, df = 9, p = 0.044) differed between pitfall trapping rounds. No significant taxon-specific relationships were identified for the interaction between sowing period and pitfall trapping round.

**Figure 1:**
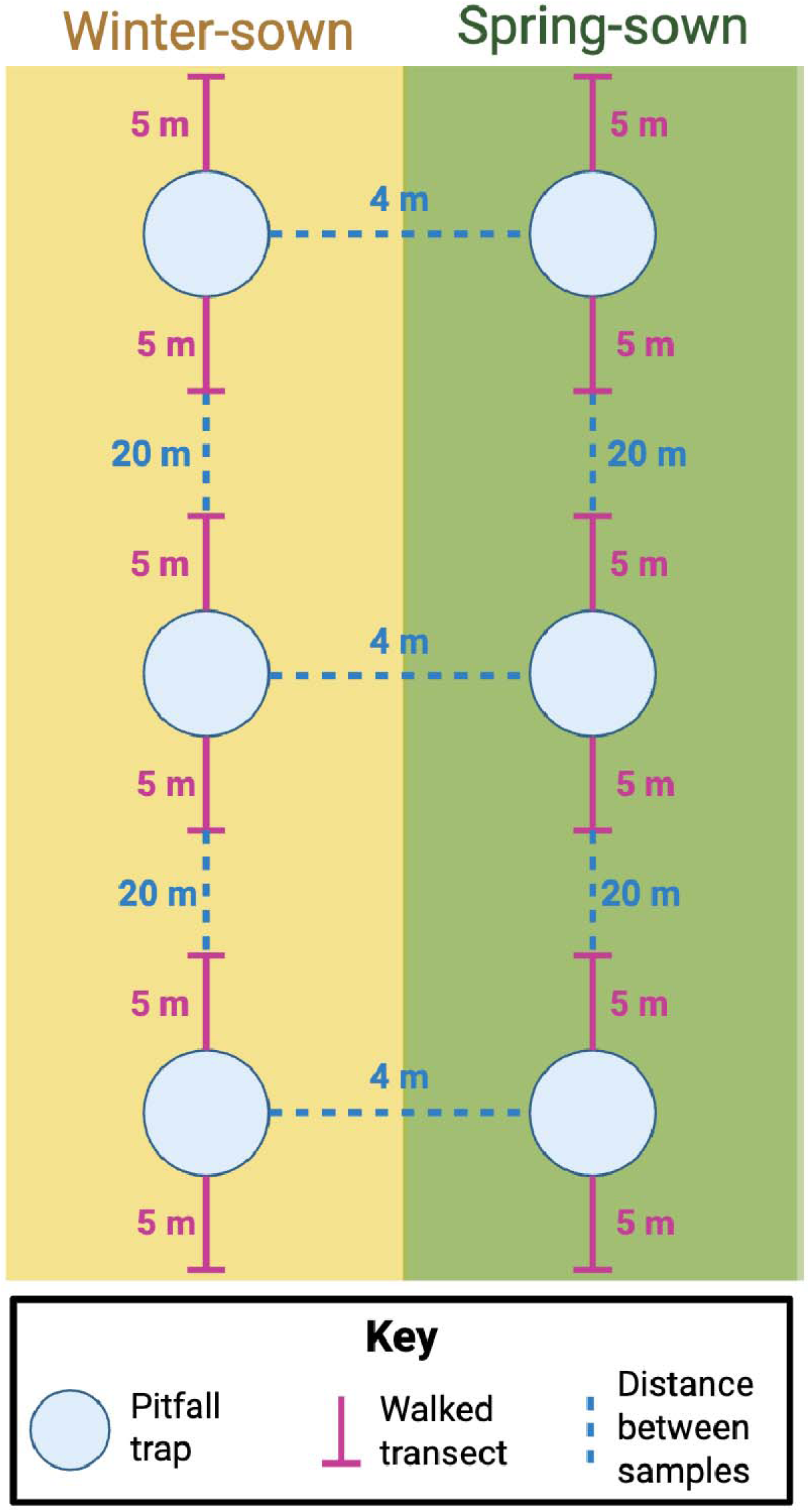
Experimental design for invertebrate sampling. Paired sampling locations were established 4 m apart in directly adjacent plots of winter- and spring-sown wheat. Created in BioRender. Cuff, J. (2025) https://BioRender.com/6rd9udt

**Figure 2:**
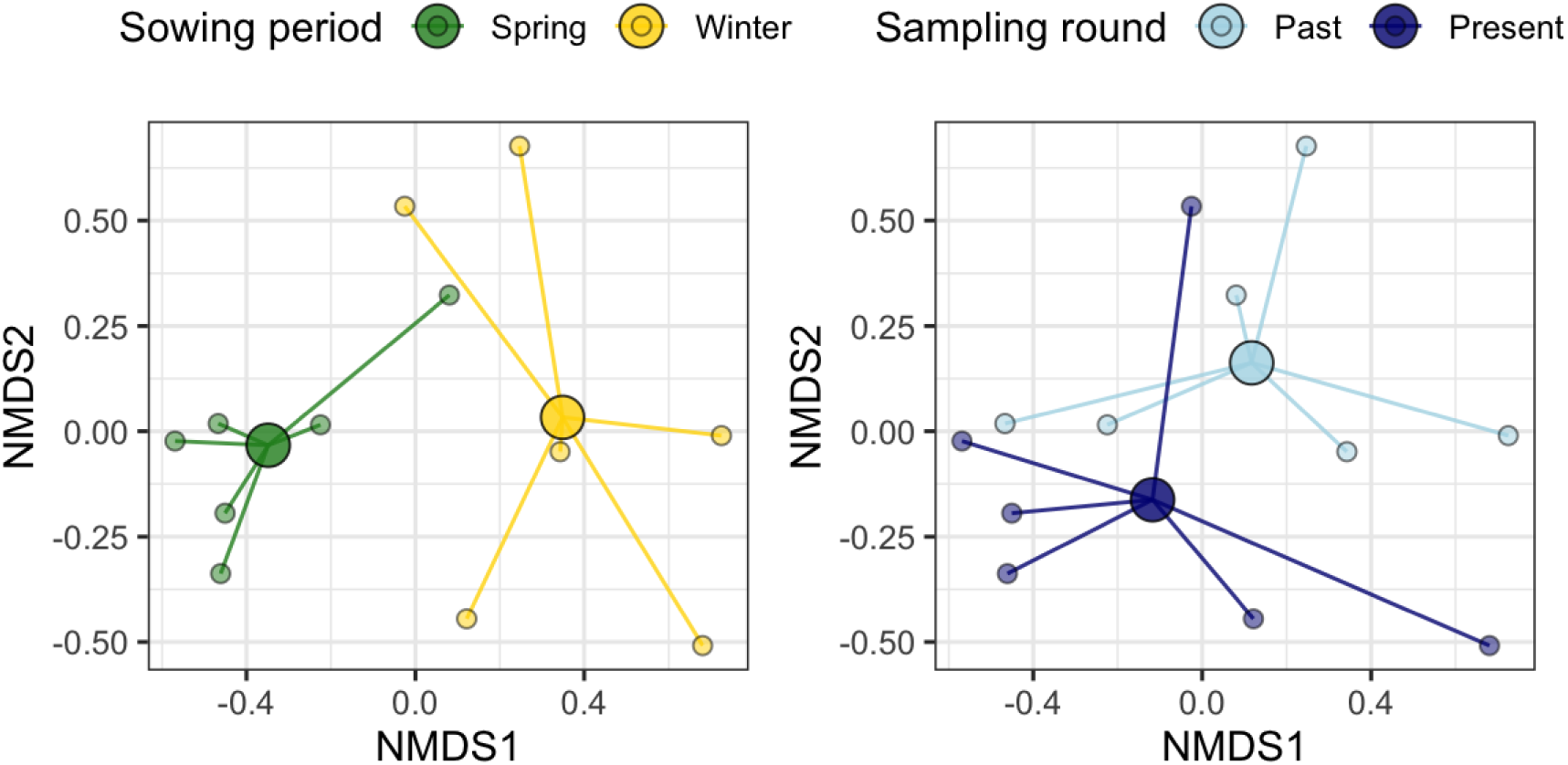
Non-metric multidimensional scaling of arthropod prey communities available in spring-sown and winter-sown wheat, and between pitfall trapping rounds. Stress = 0.103.

Predatory arthropod nutrient contents significantly differed between crops sown in winter and spring (MLM: F_1,87_ = 7.537, p = 0.002; Figure 3), but this depended on the taxon (MLM: F_9,83_ = 1.986, p = 0.002). Arthropod nutrient contents did not, however, significantly differ between taxa alone. Specifically, the proportional content of carbohydrate (MLM: F_1,87_ = 5.351, p = 0.002), lipid (MLM: F_1,87_ = 2.130, p = 0.002) and protein (MLM: F_1,87_ = 0.056, p = 0.002) significantly differed between crops sown in winter and spring, with carbohydrate being proportionally higher in winter-sown crops and lipid being proportionally higher in spring-sown crops. All three macronutrients were more variable in their content in winter-sown crops. As well, the percentage content of carbohydrate (MLM: F_9,83_ = 0.660, p = 0.002), lipid (MLM: F_9,83_ = 1.035, p = 0.002) and protein (MLM: F_9,83_ = 0.291, p = 0.002) significantly differed based on the interaction between sowing period and taxon.

**Figure 3:**
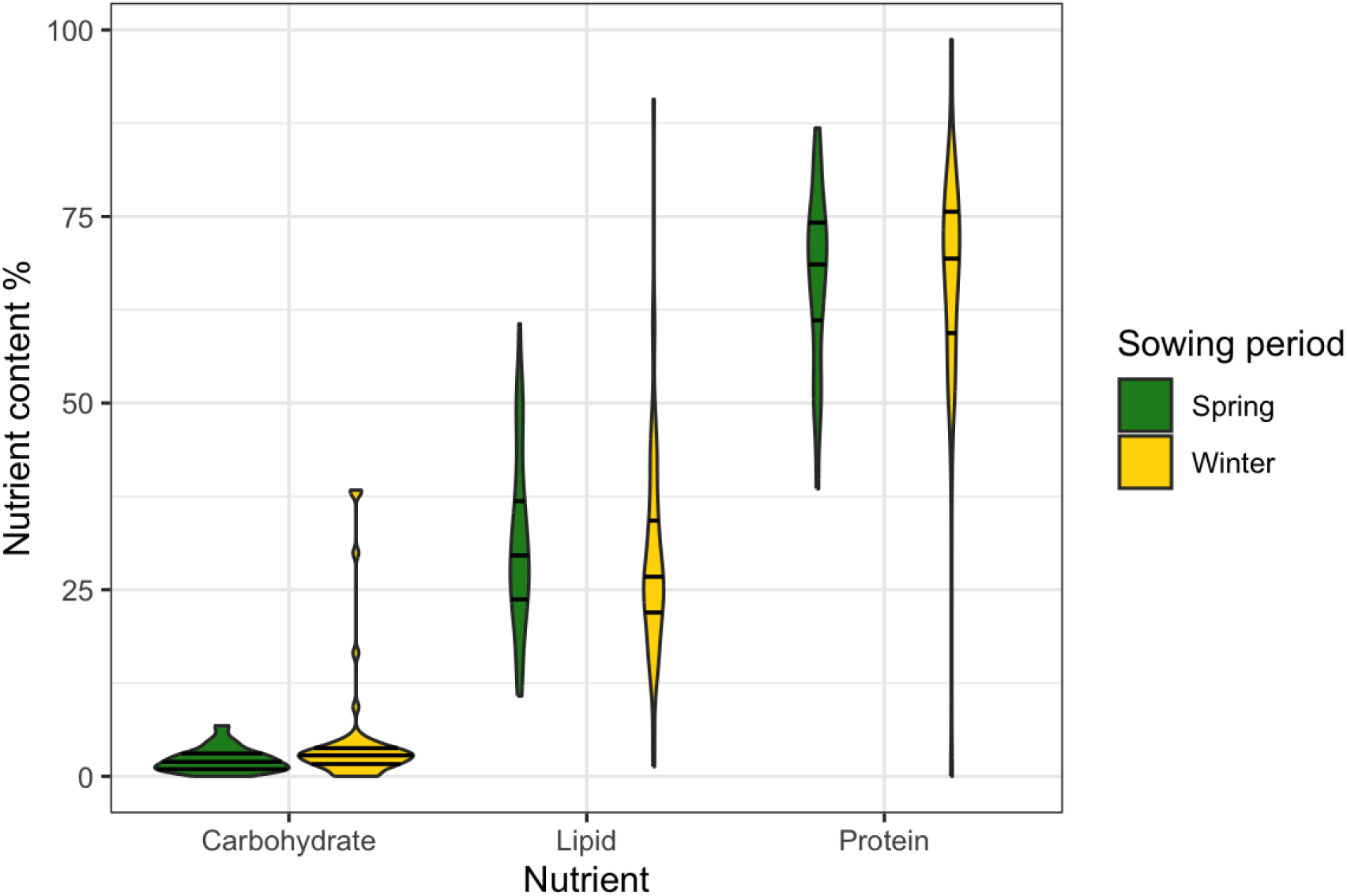
Violin plot of proportional macronutrient content (carbohydrate, lipid, protein) of predatory arthropods collected from winter-sown and spring-sown wheat. Horizontal lines within each violin represent the median and quartiles.

Null network predator prey interactions did not significantly explain predator nutrient contents when considering past (RDA: F = 0.851, df = 19, p = 0.673: Figure 4) nor present (RDA: F = 0.864, df = 24, p = 0.656: Figure 4) prey availability, and most individual model terms (prey) did not significantly explain predator nutrient content, although there was a marginally insignificant relationship between past Keroplatidae abundance and predator nutrient contents (ANOVA: F = 2.686, df = 1, p = 0.089), and significant and marginally insignificant relationships involving present Collembola (ANOVA: F = 6.361, df = 1, p = 0.013) and Braconidae (ANOVA: F = 2.662, df = 1, p = 0.096) abundances, respectively.

**Figure 4:**
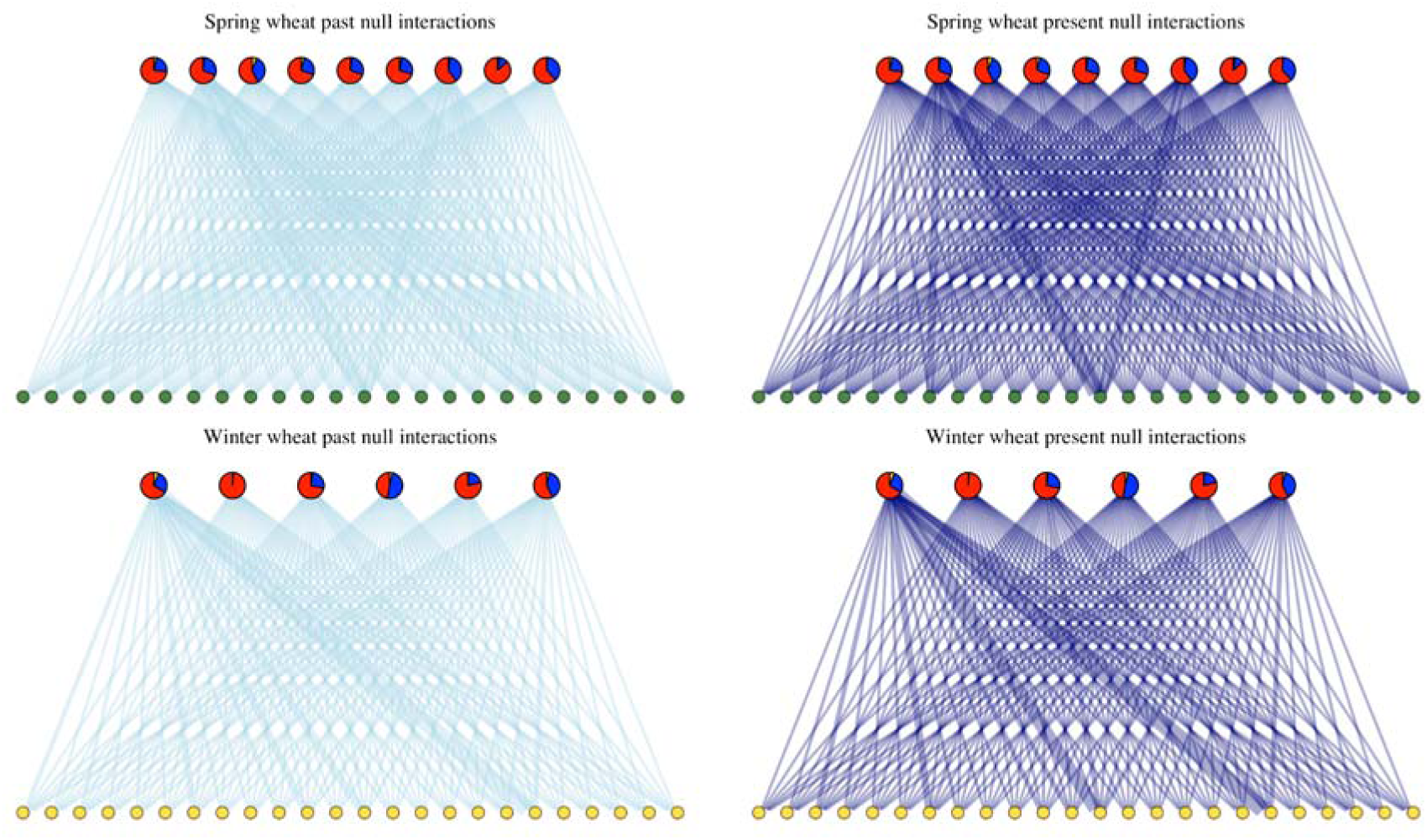
Inferred networks of predator-prey interactions based on prey availability for spring wheat (green prey nodes) and winter wheat (yellow prey nodes) based on data taken nine days before (‘past’; light blue links) and in parallel with samples used for nutritional analysis (‘present’; dark blue links). Upper nodes are predators when present nutrient data given with yellow, blue and red denoting carbohydrate, lipid, and protein proportional content, respectively.

## Discussion

The nutrient content of arthropod predators was significantly associated with the timing of crop sowing, even across the small spatial scales used in this study (i.e., parallel transects 4 m apart). Whilst the prey available to those predators also changed across those small spatial scales, neither current nor recent prey availability explained the nutritional differences. This could indicate density-independent foraging of the arthropod predators (i.e., foraging for prey disproportionate to their availability, observed in similar predatory assemblages; Cuff et al., 2022), or a shift in prey quality, irrespective of taxonomic identity, associated with differences in the developmental stage of crops sown in winter and spring. The crops themselves will likely differ markedly in their nutrient content at different developmental stages given the remobilisation of nutrients away from leaf tissue over time (Gregersen et al., 2008). This will possibly have concomitant impacts to the herbivorous insects feeding on them, which are then predated by the focal arthropod predators of this study. The greater variability in nutrient content in predators from the winter-sown wheat, for example, could relate to the scarcity of nutrients in the senescing wheat relative to the younger wheat in the spring-sown crop. Senescing crops can drive the death or dispersal of foliar herbivores and their predators (Watt, 1979), likely resulting in nutritional variation and decline in those that remain. Prey quality, rather than availability, may therefore be the primary driver of the observed nutritional differences (Rendon et al., 2019; Schmidt et al., 2012; Wilder et al., 2010). Since prey nutrient content influences the nutrition and trophic interactions of predators (Cuff, Tercel, Vaughan, et al., 2024; Wilder et al., 2019), this could indicate a nutritional cascade from the crop as a basal resource to the arthropod predators.

This hypothetical nutritional cascade has important implications for subsequent biocontrol and the spillover of arthropods and interactions from winter- to spring-sown crops continuously and following harvest. The greater nutritional variation in predators in the winter-sown wheat could indicate increased rates of nutritional imbalance, which would then be likely redressed through nutrient-specific foraging (Mayntz et al., 2005). Given the wide variation either side of the nutrient contents of spring-sown wheat predators in winter-sown wheat predators, this may confound predictions of interactions of these predators as they migrate to other locations, such as spring-sown crops. Harvest of winter-sown crops, which will increase community turnover through physical disturbance (Cuff et al., 2021; Opatovsky & Lubin, 2012), is likely to propagate this migration through spillover, especially between adjacent plots (Cuff, Gajski, et al., 2024).

This work was intentionally carried out at a small spatial scale, with only 4 m separating the transects in winter- and spring-sown crops. The intention was to determine either that differences could resolve at such small spatial scales, or that arthropods were spilling over between the two adjacent crops. Whilst a significant difference in nutrient content was detected at this resolution, spillover is likely nonetheless, especially given that many of the arthropod predators studied are mobile. Most of the predators collected were small ground beetles such as *Bembidion* spp., the foraging range of which is poorly characterised; however, other larger ground beetles are known to travel less than 1 m per day (Schreiner & Irmler, 2009). This could imply that the ‘present’ prey availability data were representative of their recent prey availability, even if the ‘past’ prey availability data were less likely to be representative. That a significant difference in nutrient content was observed supports the notion that either the difference between the two crop stages resulted in their divergent nutritional states or that their nutritional state drove them to forage in a specific crop stage. The latter would be consistent with optimal foraging theory if each ‘patch’ presented different nutritional opportunities (Pyke et al., 1977). Both eventualities do, however, indicate a nutritional effect of crop developmental stage with implications for wider foraging and biocontrol.

This study was a brief snapshot of a potentially complex and continuously evolving dynamic between crop development and predator nutrition. Given the indirect link between predators and crops, other nutrients, including elemental macronutrients and micronutrients, may be important drivers of interactions (Kaspari, 2021; Wilder et al., 2025). This study provides the foundations from which to investigate this dynamic deeper, ideally by linking nutrient flow through each trophic level to determine how nutrients structure the wider ecological system (Cuff, Evans, et al., 2024). Whilst the interactions inferred based on prey availability data are likely to reflect the pool of prey accessible to the predators studied and null models can generate semi-accurate interaction data (Cuff, Tercel, Windsor, et al., 2024), trophic interactions are often density-independent (Cuff et al., 2022; Gajski et al., 2024; Vaughan et al., 2018), the deviations of which from our models are unknown. The interactions inferred should therefore be interpreted with caution, and future investigations should identify the interactions empirically.

## Conclusions

We have demonstrated that the nutrient contents of arthropod predators can change with the sowing period of crops on even very small spatial scales. Whilst prey availability concomitantly changes, it does not explain this nutritional variation, possibly indicating a more complex set of mechanisms, potentially relating to cascading impacts to the nutritional quality of prey as a result of different stages of crop development and senescence. That basal resource quality could underpin predator nutritional fitness through trophic cascades has important implications for the management of crops to sustain healthy natural enemy populations. This is likely to be particularly relevant in monocultural arable systems where alternative basal resources are scarce if available at all. The potential spillover of nutritionally distinct predator and pest populations from winter- to spring-sown crops, either continuously or following harvest, has the potential to rewire nutritional networks in adjacent spring-sown crops. This study provides the foundations for further investigation of this crucial nutritional dynamic. Expansion of the spatiotemporal scope of this study could identify novel management strategies to optimise natural enemy population health and conservation biocontrol.

## Author contributions

Rosy Christopher: data curation; formal analysis; investigation; methodology; visualisation; writing - review and editing. Rebecca Wright: conceptualization; data curation; formal analysis; investigation; methodology; visualisation; writing – original draft. Fredric M. Windsor: investigation; methodology; project administration; supervision; writing - review and editing. Jordan P. Cuff: conceptualization; data curation; formal analysis; investigation; methodology; project administration; supervision; visualisation; writing - review and editing.

## Conflicts of interest

None to declare.

## Acknowledgments

RC and JPC were funded by a Newcastle University Academic Track Fellowship. For the purpose of open access, the author has applied a Creative Commons Attribution (CC-BY) licence to any Author Accepted Manuscript version arising from this submission.

## Data availability

All data and code are openly available via Zenodo: https://doi.org/10.5281/zenodo.17911631

## Notes

https://doi.org/10.5281/zenodo.17911631

### Competing Interest Statement

The authors have declared no competing interest.

## References

Barber, A. D. (2008). Key to the identification of British centipedes (1. ed). Field Studies Council.

Briatte, F. (2021). ggnetwork: Geometries to plot networks with ‘ggplot2’ [Computer software].

Csardi, G., & Nepusz, T. (2006). The igraph software package for complex network research. InterJournal Complex Systems, 1695, 1–9.

Cuff, J. P., Drake, L. E., Tercel, M. P. T. G., Stockdale, J. E., Orozco□terWengel, P., Bell, J. R., Vaughan, I. P., Müller, C. T., & Symondson, W. O. C. (2021). Money spider dietary choice in pre□ and post□harvest cereal crops using metabarcoding. Ecological Entomology, 46(2), 249–261. 10.1111/een.12957

Cuff, J. P., Evans, D., Vaughan, I., Wilder, S., Tercel, M., & Windsor, F. (2024). Networking nutrients: How nutrition determines the structure of ecological networks. Journal of Animal Ecology, 93(8), 974–988. 10.1111/1365-2656.14124

Cuff, J. P., Gajski, D., Michalko, R., Košulič, O., & Pekár, S. (2024). Biomonitoring of biocontrol across the full annual cycle in temperate climates: Post□harvest, winter and early□season interaction data and methodological considerations for its collection. Agricultural and Forest Entomology, afe.12635. 10.1111/afe.12635

Cuff, J. P., Tercel, M. P. T. G., Drake, L. E., Vaughan, I. P., Bell, J. R., Orozco□terWengel, P., Müller, C. T., & Symondson, W. O. C. (2022). Density□independent prey choice, taxonomy, life history, and web characteristics determine the diet and biocontrol potential of spiders (Linyphiidae and Lycosidae) in cereal crops. Environmental DNA, 4(3), 549–564. 10.1002/edn3.272

Cuff, J. P., Tercel, M. P. T. G., Vaughan, I. P., Drake, L. E., Wilder, S. M., Bell, J. R., Müller, C. T., Orozco□terWengel, P., & Symondson, W. O. C. (2024). Prey nutrient content is associated with the trophic interactions of spiders and their prey selection under field conditions. *Oikos*, e10712. 10.1111/oik.10712

Cuff, J. P., Tercel, M. P. T. G., Windsor, F. M., Hawthorne, B. S. J., Hambäck, P. A., Bell, J. R., Symondson, W. O. C., & Vaughan, I. P. (2024). Sources of prey availability data alter interpretation of outputs from prey choice null networks. Ecological Entomology, 49(3), 418–432. 10.1111/een.13315

Cuff, J. P., Windsor, F. M., Tercel, M. P. T. G., Bell, J. R., Symondson, W. O. C., & Vaughan, I. P. (2023). Temporal variation in spider trophic interactions is explained by the influence of weather on prey communities, web building and prey choice. Ecography, e06737. 10.1111/ecog.06737

Dallimore, T., & Shaw, P. J. A. (with Field Studies Council (Great Britain)). (2013). Illustrated key to the families of British springtails (Collembola). FSC Publications.

Douglas, D. J. T., Vickery, J. A., & Benton, T. G. (2010). Variation in arthropod abundance in barley under varying sowing regimes. Agriculture, Ecosystems & Environment, 135(1–2), 127–131. 10.1016/j.agee.2009.09.002

Gajski, D., Mifková, T., Košulič, O., Michálek, O., Serbina, L. Š., Michalko, R., & Pekár, S. (2024). Brace yourselves, winter is coming: The winter activity, natural diet, and prey preference of winter-active spiders on pear trees. Journal of Pest Science, 97, 113–126. 10.1007/s10340-023-01609-5

Gregersen, P. L., Holm, P. B., & Krupinska, K. (2008). Leaf senescence and nutrient remobilisation in barley and wheat. Plant Biology, 10(s1), 37–49. 10.1111/j.1438-8677.2008.00114.x

Hedlund, K., Griffiths, B., Christensen, S., Scheu, S., Setälä, H., Tscharntke, T., & Verhoef, H. (2004). Trophic interactions in changing landscapes: Responses of soil food webs. Basic and Applied Ecology, 5(6), 495–503. 10.1016/j.baae.2004.09.002

Heimbach, U., & Garbe, V. (1996). Effects of reduced tillage systems in sugar beet on predatory and pest arthropods. Acta Jutlandica, 71(2), 195–208.

Hodge, M. A. (1999). The Implications of Intraguild Predation for the Role of Spiders in Biological Control. The Journal of Arachnology, 27(1), 351–362.

Hsu, G., Ou, J., & Ho, C. (2021). Pest consumption by generalist arthropod predators increases with crop stage in both organic and conventional farms. Ecosphere, 12(7), e03625. 10.1002/ecs2.3625

Jaworski, C. C., Thomine, E., Rusch, A., Lavoir, A.-V., Wang, S., & Desneux, N. (2023). Crop diversification to promote arthropod pest management: A review. Agriculture Communications, 1(1), 100004. 10.1016/j.agrcom.2023.100004

Jensen, K., Mayntz, D., Toft, S., Clissold, F. J., Hunt, J., Raubenheimer, D., & Simpson, S. J. (2012). Optimal foraging for specific nutrients in predatory beetles. Proceedings of the Royal Society B: Biological Sciences, 279(1736), 2212–2218. 10.1098/rspb.2011.2410

Jowett, K., Milne, A. E., Garrett, D., Potts, S. G., Senapathi, D., & Storkey, J. (2021). Above□ and below□ground assessment of carabid community responses to crop type and tillage. Agricultural and Forest Entomology, 23(1), 1–12. 10.1111/afe.12397

Kaspari, M. (2021). The Invisible Hand of the Periodic Table: How Micronutrients Shape Ecology. Annual Review of Ecology, Evolution, and Systematics, 52(1), 199–219. 10.1146/annurev-ecolsys-012021-090118

Landa, B. B., Navas-Cortés, J. A., & Jiménez-Díaz, R. M. (2004). Integrated Management of Fusarium Wilt of Chickpea with Sowing Date, Host Resistance, and Biological Control. Phytopathology®, 94(9), 946–960. 10.1094/PHYTO.2004.94.9.946

Luff, M. L. (2007). Handbooks for the identification of British insects*. Vol.* 4*, Pt. 2*: *The Carabidae (ground beetles) of Britain and Ireland* (2. ed). Royal Entomological Society.

Mayntz, D., Raubenheimer, D., Salomon, M., Toft, S., & Simpson, S. J. (2005). Nutrient-Specific Foraging in Invertebrate Predators. Science, 307(5706), 111–113. 10.1126/science.1105493

Michalko, R., Pekár, S., Dul’a, M., & Entling, M. H. (2019). Global patterns in the biocontrol efficacy of spiders: A meta□analysis. Global Ecology and Biogeography, 28(9), 1366–1378. 10.1111/geb.12927

Moore, S. M., Borer, E. T., & Hosseini, P. R. (2010). Predators indirectly control vector-borne disease: Linking predator-prey and host-pathogen models. Journal of the Royal Society, Interface, 7(42), 161–176. 10.1098/rsif.2009.0131

Oksanen, J., Simpson, G. L., Blanchet, F. G., Kindt, R., Legendre, P., Minchin, P. R., O’Hara, R. B., Solymos, P., Stevens, M. H. H., Szoecs, E., Wagner, H., Barbour, M., Bedward, M., Bolker, B., Borcard, D., Carvalho, G., Chirico, M., De Caceres, M., Durand, S., … Weedon, J. (2022). vegan: Community Ecology Package (Version 2.6-2) [Computer software]. https://CRAN.R-project.org/package=vegan

Opatovsky, I., & Lubin, Y. (2012). Coping with abrupt decline in habitat quality: Effects of harvest on spider abundance and movement. Acta Oecologica, 41, 14–19. 10.1016/j.actao.2012.03.001

Pyke, G. H., Pulliam, H. R., & Charnov, E. L. (1977). Optimal foraging: A selective review of theory and tests. The Quarterly Review of Biology, 52, 137–154.

R Core Team. (2025). R: A Language and Environment for Statistical Computing (Version 4.5.2) [Computer software]. R Foundation for Statistical Computing. https://www.R-project.org/

Rendon, D., Taylor, P. W., Wilder, S. M., & Whitehouse, M. E. A. (2019). Does prey encounter and nutrient content affect prey selection in wolf spiders inhabiting Bt cotton fields? PLOS ONE, 14(1), e0210296. 10.1371/journal.pone.0210296

Roberts, M. J. (2016). Spiders of Britain and Northern Europe (Reprint).

Rosenblatt, A. E. (2018). Shifts in plant nutrient content in combined warming and drought scenarios may alter reproductive fitness across trophic levels. Oikos, 127(12), 1853–1862. 10.1111/oik.05272

Rosenheim, J. A., Kaya, H. K., Ehler, L. E., Marois, J. J., & Jaffee, B. A. (1995). Intraguild Predation Among Biological-Control Agents: Theory and Evidence. Biological Control, 5(3), 303–335. 10.1006/bcon.1995.1038

Schmidt, J. M., Sebastian, P., Wilder, S. M., & Rypstra, A. L. (2012). The Nutritional Content of Prey Affects the Foraging of a Generalist Arthropod Predator. PLoS ONE, 7(11), e49223. 10.1371/journal.pone.0049223

Schreiner, R., & Irmler, U. (2009). Mobility and Spatial use of the Ground Beetle species Elaphrus cupreus and Elaphrus uliginosus (Coleoptera: Carabidae). Entomologia Generalis, 32(3), 165–179.

Symondson, W. O. C., Sunderland, K. D., & Greenstone, M. H. (2002). Can Generalist Predators be Effective Biocontrol Agents? Annual Review of Entomology, 47(1), 561–594. 10.1146/annurev.ento.47.091201.145240

Tilling, S. (2014). A key to the major groups of British terrestrial invertebrates (Second edition). FSC.

Tschumi, M., Ekroos, J., Hjort, C., Smith, H. G., & Birkhofer, K. (2018). Predation□mediated ecosystem services and disservices in agricultural landscapes. Ecological Applications, 28(8), 2109–2118. 10.1002/eap.1799

Vaughan, I. P., Gotelli, N. J., Memmott, J., Pearson, C. E., Woodward, G., & Symondson, W. O. C. (2018). econullnetr: An _R_ package using null models to analyse the structure of ecological networks and identify resource selection. Methods in Ecology and Evolution, 9(3), 728–733. 10.1111/2041-210X.12907

Wang, Y., Naumann, U., Wright, S. T., & Warton, D. I. (2012). Mvabund– an R package for model□based analysis of multivariate abundance data. Methods in Ecology and Evolution, 3(3), 471–474. 10.1111/j.2041-210X.2012.00190.x

Watt, A. D. (1979). The effect of cereal growth stages on the reproductive activity of Sitobion avenue and Metopolophium dirhodum. Annals of Applied Biology, 91(2), 147–157. 10.1111/j.1744-7348.1979.tb06485.x

Wickham, H. (2016). ggplot2: Elegant Graphics for Data Analysis (2nd ed. 2016). Springer International Publishing□: Imprint: Springer. 10.1007/978-3-319-24277-4

Wickham, H., Averick, M., Bryan, J., Chang, W., McGowan, L., François, R., Grolemund, G., Hayes, A., Henry, L., Hester, J., Kuhn, M., Pedersen, T., Miller, E., Bache, S., Müller, K., Ooms, J., Robinson, D., Seidel, D., Spinu, V., … Yutani, H. (2019). Welcome to the Tidyverse. Journal of Open Source Software, 4(43), 1686. 10.21105/joss.01686

Wilder, S. M., Barnes, C. L., & Hawlena, D. (2019). Predicting Predator Nutrient Intake From Prey Body Contents. Frontiers in Ecology and Evolution, 7, 42. 10.3389/fevo.2019.00042

Wilder, S. M., Herzog, C., Reeves, J., Knowles, O., & Cuff, J. P. (2025). Bridging digestive physiology and ecology for a more integrative understanding of invertebrate predators. Journal of Experimental Biology, 228(14), jeb249697. 10.1242/jeb.249697

Wilder, S. M., Mayntz, D., Toft, S., Rypstra, A. L., Pilati, A., & Vanni, M. J. (2010). Intraspecific variation in prey quality: A comparison of nutrient presence in prey and nutrient extraction by predators. Oikos, 119(2), 350–358. 10.1111/j.1600-0706.2009.17819.x

